# Small heat shock proteins (sHSPs) identified in nodules of tropical woody legumes

**DOI:** 10.1101/2024.10.07.617047

**Authors:** Cara G. Flynn, Rayan Fakih, Kalle Gehring, Fiona M. Soper

**Affiliations:** Department of Biochemistry, McGill University, Montreal, QC, Canada; Department of Biology, McGill University, Montreal, QC, Canada; Bieler School of Environment, McGill University, Montreal, QC, Canada

**Keywords:** *Acacia*, biogeochemical cycling, Fabaceae, Hsp, Nitrogen fixation, *Rhizobia*, warming

## Abstract

Biological nitrogen fixation (BNF) is a primary input of nitrogen to natural and agricultural systems globally. BNF is a temperature-dependant enzymatic process and can be conducted by microbes (including *Rhizobia*) hosted symbiotically in root nodules of some plants. Heat shock proteins (Hsps) have been implicated in the process of acquiring thermotolerance or acclimating to elevated temperature, as they play a vital role in maintaining cell integrity and homeostasis during heat stress. Although the BNF response to temperature may crucially impact future ecosystem productivity in the face of global climate change, little is known about Hsp expression in nodules of N-fixing non-agricultural species, such as tropical N-fixing trees in the *Fabaceae* family. This project aimed to characterize small (15-20 kDa) Hsp (sHsp) expression in nodule tissue to examine the biochemical mechanisms of heat response in these tissues. To first identify Hsps in nodule tissues, *Vachellia farnesiana* and *Acacia confusa* nodules were excised, heat shock was induced, and protein content was isolated via chemical treatment before separation of protein species and analysis with SDS-PAGE. Two polyacrylamide gels yielded bands in the 15-20 kDa region that displayed differential Coomassie staining, which were sent for further characterization by HPLC-MS analysis for protein sequencing. Ten rhizobial sHsps were detected in these samples in addition to seven *Acacia* sHsps when compared independently to reference rhizobial and plant proteome databases. In an attempt quantify relative expression of Hsps in nodule and root tissue, we performed western blot experiments using Anti-Hsp20 antibodies raised against human and mouse Hsp proteins, with anti-beta actin loading control. While nonuniform beta-actin expression across tissue types (*A. confusa* nodules versus root control) prevented quantitative analysis, the experiments validated that Hsp20s are expressed in *Acacia* nodules as well as in root tissue. These experiments provide a foundation for future studies to determine variation in responses to key stressors predicted to increase with global climate change and help determine the implications of warming across the tropics and beyond. Proteomics data are available via ProteomeXchange with identifier PXD055599.

## Introduction

Climate change continues to dramatically alter abiotic processes that impact the biosphere and there remain many unanswered questions about the effect of warming on key drivers of ecosystem function, such as biological nitrogen fixation (BNF). As one of the primary elements required to sustain the structure and function of living organisms, the cycling of nitrogen (N) between the atmosphere, lithosphere, and biosphere is of great concern to ecosystem scientists. Despite high rates of N inputs to agricultural systems via chemical fertilization, BNF remains the primary N input in natural ecosystems, contributing an estimated minimum of 100 Tg of N annually in both agricultural and natural ecosystems^1^. BNF is the mechanism by which prokaryotes convert inorganic, atmospheric N_2_ into a bioavailable form. Often these microbes are symbionts which rely on a host plant to perform N-fixation (e.g., rhizobia)^2^. Symbiotic N-fixers are of particular interest as they exhibit the greatest rates of N-fixation^2^. To host their symbiont, plants develop structures known as nodules that facilitate microanoxic environments conducive to N-fixation in which rhizobia differentiate into N-fixing bacterioids^3^. Nitrogenase is the prokaryotic enzyme that converts N_2_ into ammonia (NH_4_^+^), a bioavailable form that can be assimilated into plant metabolism and used for essential biochemical processes like amino acid and nucleic acid synthesis^3^. In exchange for fixed N, plants tightly control oxygen in the nodule tissues and provide carbon (C) energy sources to support bacterial metabolism^3,4^. This nuanced interaction has been observed to respond to a variety of abiotic environmental factors, and N-fixation is seen to be regulated at multiple scales including investments in nodule construction – changes in the quantity and/or total biomass of nodules – or plant physiological responses which impact measured rates of fixation in nodule tissue^5^.

Considering BNF is an enzymatic process that will presumably be affected by the increasing air and soil temperatures observed globally in recent years, ongoing climate change is likely to impact this and other N-cycling processes, which in turn alter global food security, net primary productivity (NPP), and ecosystem function. Recent studies investigating BNF in woody plant species have revealed that fixation rates may vary more than previously estimated across temperature ranges and biomes, with tropical species reaching temperature optima as high as 36 ºC, far hotter than the previously estimated and widely cited value of 25 ºC^5^. Moreover, it was recently observed that the relationship between BNF and temperature was best modeled by an asymmetric function, resembling a “tipping point” dynamic, which is consistent with enzymatic temperature response curves^5^, rather than the previously assumed symmetric relationship depicting a more gradual downregulation response^6^. Current terrestrial biosphere models (TBMs) are often operating with outdated N input and acquisition functions that do not account for these parameters nor do they include acclimation potential, despite research suggesting that species are capable of acclimating to elevated temperatures^5^. Incorporating estimates of BNF variability and enhancing our mechanistic understanding of the relationship between BNF and temperature is essential for improving our predictions of future ecosystem C storage capabilities, which have implications for conservation practices, climate goals, and mitigation strategies. Understanding the biochemical mechanisms that underlie N-fixation regulation and acclimation of N-fixation to elevated temperatures will improve our predictions as to how BNF will respond to global climate change.

Heat shock response (HSR) is a well-known eukaryotic biochemical response to temperature that has been observed in a variety of plant tissues. Exposure to heat stress, or exposure to temperatures above a threshold – typically 10-15 ºC above optimal growing conditions – can result in abnormal increases in membrane fluidity, degradation, inactivation of enzymes, and often a toxic aggregation of proteins in cells^7^. Heat Shock Response is often characterized by the onset of Heat Shock Protein (Hsp) expression which preserves cell resilience during heat stress by facilitating protein folding and downregulating global transcription to prevent protein aggregation in cells^7^. Hsps are upregulated during and shortly after cell exposure to heat stress and are classified primarily by size and function^8^. Heat stress can vary in the degree of temperature change and duration of exposure; in natural ecosystems, this is dependent on the probability that a climatic zone will reach and maintain elevated temperatures^7^. Hsp expression has been observed across a taxonomically diverse range of organisms, from eukaryotic groups such as mammals and insects to some prokaryotes, suggesting the HSR is a universal mechanism for combatting heat stress^7^. Hsp expression has been observed in all types of plant tissues, including leaf, flower, fruit, and root tissues, and an Hsp has even been implicated in the nodulation process in soybeans^9^. Hsps are classified according to their approximate molecular weight which can range from 10 to 200 kDa, and occasionally are classified by amino acid sequence and homology^8^. Five Hsp classes have been described for plants: Hsp100 family, Hsp90 family, Hsp70 family, Hsp60 family, and small Hsp (sHsp) family^8^. Plants occupy an extremely wide range of climatic conditions and must express traits on both short and long-term time scales in order to maximize a specific species’ ability to thrive in any given place at a particular time^10^. Plants are sessile organisms which rely on environmental cues as a result of immobility, and therefore rely on acclimation, e.g., acquired thermotolerance, to adapt to harsh environments and survive otherwise lethal temperatures^7^. In a proteomic analysis of sHsp expression in a hybrid thermotolerant maize relative to its thermosensitive parent, it was observed that 22 and 27 kDa Hsps were overexpressed in the thermotolerant strain, indicating these sHsps could play a major role in acquired thermotolerance^11^. This acclimation response is of particular importance to long-lived woody plant species as they may be exposed to temperature stressors that can compound or become exacerbated over time, and their population dynamics occur over longer periods of time, discounting range shifts or genetic adaptations as viable methods for short-term stress avoidance. Despite the wide scope of literature that implicates Hsps as a mechanism of acquired thermotolerance and acclimation in many plant tissues, and although temperature has been demonstrated to be a crucial regulator of BNF^5^, characterization of Hsp expression in tropical N-fixing tree nodules has yet to be examined as a mechanism of regulation during heat stress and acclimation.

The primary objective of this project was to determine whether Hsps, specifically sHsps, are expressed in tropical legume tree nodules and play a role in the thermoacclimation observed in previous N-fixation studies^5^. Our secondary aim was to quantify relative sHsp expression across species and tissue types (fine root vs nodule tissues), to investigate whether Hsp expression may be a mechanism used to maintain cellular homeostasis in nodules to facilitate N-fixation at elevated temperatures. We predicted that tropical legume species would express more sHsps in response to heat shock than temperate species, if Hsp expression is indeed a mechanism of thermoacclimation driving differential optima temperature in N-fixation across these groups of N-fixers. Furthermore, we would expect both plant and bacterial sHsps will be expressed in nodule tissues, as both cell types compose these plant organs. However, it is unknown in what proportion these species will express Hsps and whether or not root tissues will display differential expression relative to nodule extracts which may further indicate if one specific symbiotic partner drives the Heat Shock Response in nodule tissues.

## Materials and Methods

### Growth conditions and inoculation of tropical woody legumes

Seeds of Fabaceae species *Vachellia farnesiana* (formerly *Acacia farnesiana*), *Acacia confusa* (both trees native to the tropics and subtropics), and *Robinia pseudoacacia* (tree with a broad introduced distribution from tropical to temperate zones) were supplied by Sheffield’s Seed Company (NY, USA). Seeds were planted in 50:50 black earth:sand mix (v:v) and germinated in a greenhouse set at 26ºC/20ºC day/night temperature with a 16 h photoperiod of at least 600 μmols m^-2^ s^-1^ light intensity. Seedlings were transplanted to tall tree pots and inoculated after four weeks with a mixture of 11 strains of cultured *Rhizobia* and *Bradyrhizobia* in liquid media that are known to associate with *Acacia* and closely related species *in situ* (Table S1, USDA National Rhizobial Germplasm Collection). Individuals were fertilized once per week with 50 mL of a modified Hoagland’s recipe (Table S2) with a limiting quantity of N to encourage nodule formation and N-fixation in nodules.

### Heat shock induction of treated nodules and roots

Seven-to ten-week-old plants (0.5-1 m tall) were separated into treatment and control groups. In SDS-PAGE/HPLC-MS experiments, nodules were excised with root segments attached, nodules were wrapped in wet paper wipes to prevent desiccation and sealed in a glass scintillation vial. Heat-treated samples were exposed to 45 ºC heat treatment in a drying oven (Thermofisher Heratherm) for 2-3 hours while control samples were left at room temperature (14-17 ºC), with limited natural light exposure for the duration of heat treatment. For western blot experiments, whole-plants were subjected to the heat-treatment specified above while ‘total control’ samples were collected from an undisturbed plant in the greenhouse and immediately prepared for protein extraction. An additional ‘stress control’ was incorporated for whole-plant experiments in which a plant was removed from the greenhouse and left at room temperature with no natural or artificial light exposure for the duration of heat treatment to mimic the dark conditions within the drying oven without inducing heat-shock.

### Nodule and root protein extraction

Protein extraction protocol was adapted from Wang et al. (2006), a methodology for extracting total protein content from recalcitrant plant tissues^12^. For whole-plant experiments, immediately after heat treatment, nodule and root tissues were excised, rinsed thoroughly, suspended in liquid nitrogen and ground into a fine powder with a mortar and pestle. 0.1 to 0.3 g of tissue was transferred to a 2.0 mL Eppendorf microtube. Tissue was suspended in 10% TCA in acetone and the lysate was pelleted via microcentrifugation (IEC Micromax RF Refrigerated Microcentrifuge) at 4 ºC, 16,000 G for 3 minutes. The pelleted lysate was resuspended in 0.1 M ammonium acetate in 80% methanol, vortexed, and pelleted via microcentrifugation (4 ºC; 16,000 G; 3 min). The pelleted lysate was resuspended in 80% acetone (vortexed and centrifuged at 4 ºC; 16,000 G; 3 min). The pelleted lysate was air-dried (~15 min) and suspended in 1:1 phenol/SDS buffer (v:v, components of SDS buffer in Table S3), incubated at room temperature for 5 minutes, and centrifuged (room temperature.; 16,000 G; 3 min). The upper phase of the supernatant (~1 mL) was transferred to a clean 2 mL microtube. 0.1 M ammonium acetate in 80% MeOH was added to upper phase to precipitate protein. The protein was then pelleted via centrifugation (4 ºC; 16,000 G; 3 min). Protein pellets were washed (suspended, vortexed, and centrifuged) once with 100% methanol, followed by a wash with 80% acetone. Pellets were then air-dried (~20 min) and stored at 4 ºC before resuspension in rehydration buffer (8 M Urea, 4 mM DTT, 4% Chaps). From the recommendation of Rodrigues et al. sonification (3 cycles for 20 min at 50-60 ºC) of samples was conducted to increase protein solubilization^13^.

### Determining protein yield and concentration

Protein concentration in rehydrated samples was determined using a Bradford assay. In the Bradford assay, 20 μL of bovine serum albumin (BSA) solution of known protein concentration (0.2-1.0 mg/mL) were incubated in triplicate with 1 mL Bradford Reagent (BIORAD) for 5 minutes before measuring absorbance at 595 nm with a spectrophotometer^14^. Concentrations of plant protein extracts were compared to a standard curve of reference BSA solutions. Protein concentration in solution were used to standardize SDS-PAGE loading to ensure equal protein content transfer in Western Blots.

### SDS-PAGE and band analysis

After rehydration and determination of protein concentration, samples were resolved on a 12.5-15% Tris-Glycine SDS-polyacrylamide gel via SDS-PAGE and stained with Coomassie Brilliant Blue^15^. Protein ladder (BIORAD Dual-Colored Prestained Protein Standard) was used to assess protein migration and identify the molecular weight of potential bands of interest. Coomassie staining was imaged with BIORAD ChemiDoc Imager.

### Band digest and HPLC-MS Analysis

Protein extracts from one heat treated and one control sample of both A. confusa and V. farnesiana displayed differential Coomassie signal intensity in bands in the 15-20 ka region of the SDS-PAGE. Bands of interest were excised from 15% polyacrylamide gels using a razor blade along with a respective band in the control lane of approximately equal size. For each band, proteins were loaded onto a single stacking gel band to remove lipids, detergents, and salts. The gel band was reduced with DTT, alkylated with iodoacetic acid, and digested with trypsin. Extracted peptides were re-solubilized in 0.1% aqueous formic acid and loaded onto a Thermo Acclaim Pepmap (Thermo, 75 uM ID X 2 cm C18 3 uM beads) precolumn and then onto an Acclaim Pepmap Easyspray (Thermo, 75 uM × 15 cm with 2uM C18 beads) analytical column separation using a Dionex Ultimate 3000 uHPLC at 250 nl/min with a gradient of 2-35% organic (0.1% formic acid in acetonitrile) over 3 hours. Peptides were analyzed using a Thermo Orbitrap Fusion mass spectrometer operating at 120,000 resolution (FWHM in MS1) with HCD sequencing (15,000 resolution) at top speed for all peptides with a charge of 2+ or greater. The raw data were converted into *.mgf format (Mascot generic format) for searching using the Mascot 2.6.2 search engine (Matrix Science) against 17 databases: Acacia UniProtKB; Bradyrhizobium diazoefficiens UniProtKB; Sinorhizobium UniProtKB; Prosopsis cineraria Heat shock protein, Cluster ID: UniRefA0A2I4JFP8_9FABA; Bradyrhizobium guangdongense Hsp20 protein sequences, Accession Numbers A0A410V2F1, A0A410V3K3, and A0A410VH34; Mesorhizobium australicum proteome, UniProt Proteome ID: UP000010998; Mesorhizobium sp. LSHC420B00 proteome, UniProt Proteome ID: UP000018524; Mesorhizobium australicum proteome, UniProt Proteome ID: UP000193083; Rhizobium favelukesii proteome, UniProt Proteome ID: UP000019443; Rhizobium leguminosarum proteome, UniProt Proteome ID: UP000092691; Rhizobium lusitanum proteome, UniProt Proteome ID: UP000199205; Rhizobium sullae (Rhizobium hedysari) proteome, UniProt Proteome ID: UP000294576; Burkholderia plantarii proteome, UniProt Proteome ID: UP000031838; Burkholderia ubonensis proteome, UniProt Proteome ID: UP000065504; and Acacia pycnantha Genome, BioProject ID: PRJNA752212 (UniProt 2023). The database search results were loaded onto Scaffold Q+ Scaffold_5.0.1 (Proteome Sciences) for statistical treatment and data visualization.

### Western blot analysis

A ClustalOmega multiple sequence alignment was conducted to predict the viability of these antibodies in targeting bacterial and/or plant Hsp20s (Supplementary: Figure S1) which indicated relatively high homology across species. We also conducted preliminary experiments using rabbit polyclonal anti-Hsp70/HSPA1A antibodies (AF1663, R&D Biosystems) to test the viability of antibodies raised against non-bacterial nor plant species in these experiments with positive results (data not presented). Protein extracts (20-30µg) were resolved on a 12.5-15% polyacrylamide gel and transferred to polyvinylidene difluoride (PVDF) membrane. To confirm successful protein transfer, the membrane was stained with Ponceau solution. Nonspecific binding of antibodies was blocked with 5% BSA in TBST, then membranes were cut into sections and incubated overnight at 4 ºC with appropriate primary antibodies: recombinant rabbit monoclonal anti-Hsp20/HspB6 (MA5-38182, Invitrogen; Thermofisher Scientific) and rabbit polyclonal anti-beta-actin (A2066, Sigma-Aldrich; Millipore Sigma) (diluted 1:500, 1:1000 in TBST, respectively). Antibody complexes were detected using HRP-conjugated goat polyclonal anti-rabbit IgG secondary antibodies (111-035-046, Jackson ImmunoResearch) (diluted 1:5000 in TBST). ECL Prime Western Blotting Detection Reagent (Cytiva) was used to develop the chemiluminescent signal and membranes were imaged with BIORAD ChemiDoc Imager using 30s chemiluminescence exposure.

### BLAST protein characterization

Hypothetical protein sequences were described by McLay et al. and available on NCBI GenBank (BioProject Number: PRJNA752212)^16^. To identify protein species, the unique accession number for each protein species identified in data visualization was used to conduct Blastp analysis with a standard database search set of non-redundant protein sequences (nr). The algorithm selected was blastp (protein-protein BLAST).

## Results

### Small heat shock protein Hsp20 family protein and others observed in nodule protein extracts

HPLC-MS analysis revealed ten small heat shock proteins were present in *V. farnesiana* and *A. confusa* nodule tissue protein extracts corresponding to rhizobial Hsps in reference databases, which are displayed in Table 1. Additionally, seven plant sHsps were identified using the full *A. pycnatha* genome with hypothetical proteomic sequences for reference, with BLAST identification of protein species^16^, with BLAST details and putative Hsp for all hits compiled in Table 2. We observed a variety of small heat shock proteins (15-20 kDa) from a variety of symbiont partners, or rhizobial species, in all samples (control and heat-treated). Interestingly, bacterial Hsp hits primarily grouped into two main categories of sHsp/chaperone protein – Hsp20s and variant sHsps and bacterial GroES chaperones.

**Table 1:**
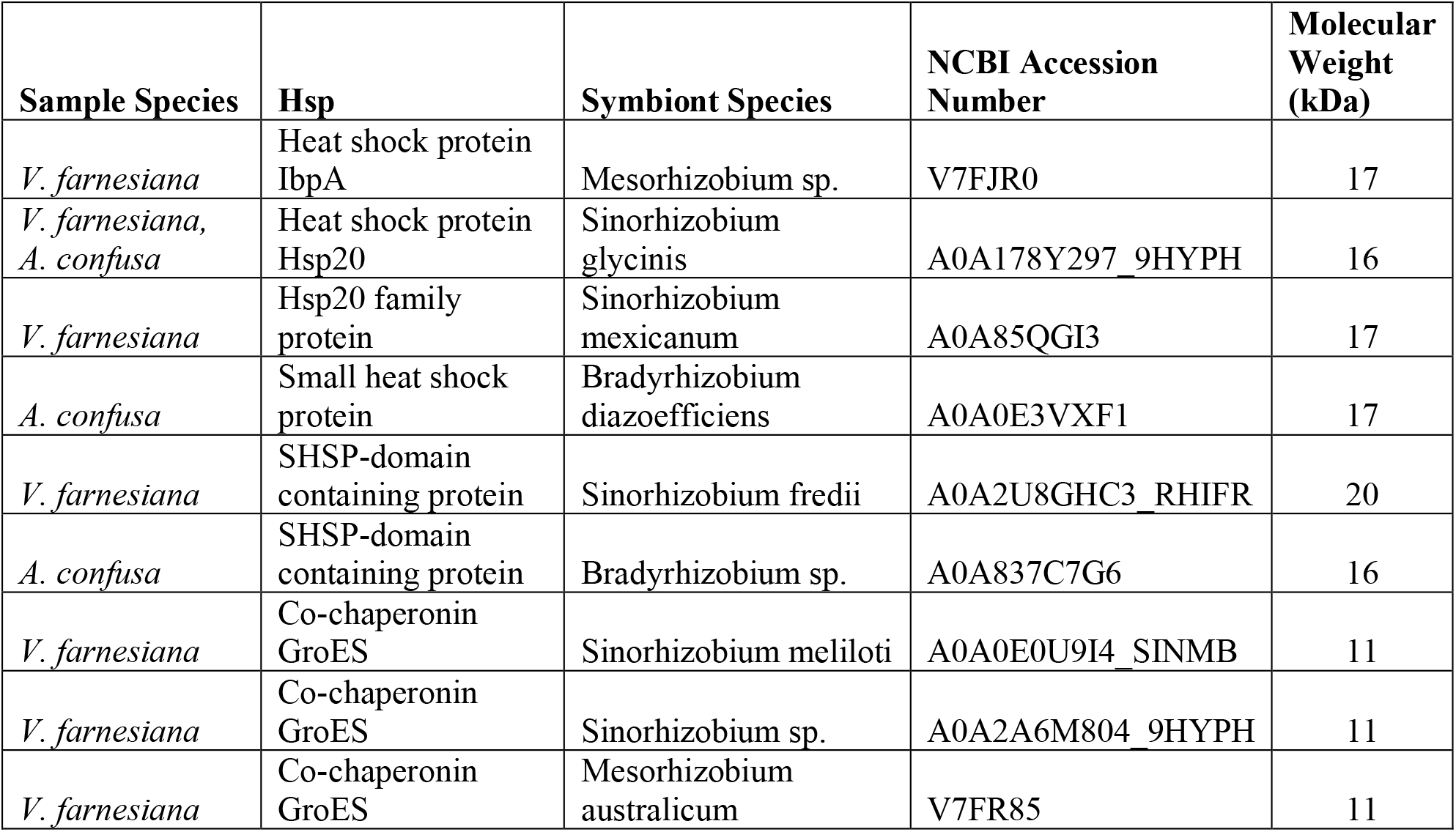
Rhizobial small heat shock proteins identified in *A. confusa* and *V. farnesiana* nodule protein extracts, characterized with HPLC-MS peptide sequencing with reference protein sequences.

| Sample Species | Hsp | Symbiont Species | NCBI Accession Number | Molecular Weight (kDa) |
| --- | --- | --- | --- | --- |
| *V. farnesiana* | Heat shock protein IbpA | Mesorhizobium sp. | V7FJR0 | 17 |
| *V. farnesiana*,<br>*A. confusa* | Heat shock protein Hsp20 | Sinorhizobium glycinis | A0A178Y297_9HYPH | 16 |
| *V. farnesiana* | Hsp20 family protein | Sinorhizobium mexicanum | A0A85QGI3 | 17 |
| *A. confusa* | Small heat shock protein | Bradyrhizobium diazoefficiens | A0A0E3VXF1 | 17 |
| *V. farnesiana* | SHSP-domain containing protein | Sinorhizobium fredii | A0A2U8GHC3_RHIFR | 20 |
| *A. confusa* | SHSP-domain containing protein | Bradyrhizobium sp. | A0A837C7G6 | 16 |
| *V. farnesiana* | Co-chaperonin GroES | Sinorhizobium meliloti | A0A0E0U9I4_SINMB | 11 |
| *V. farnesiana* | Co-chaperonin GroES | Sinorhizobium sp. | A0A2A6M804_9HYPH | 11 |
| *V. farnesiana* | Co-chaperonin GroES | Mesorhizobium australicum | V7FR85 | 11 |

**Table 2:**
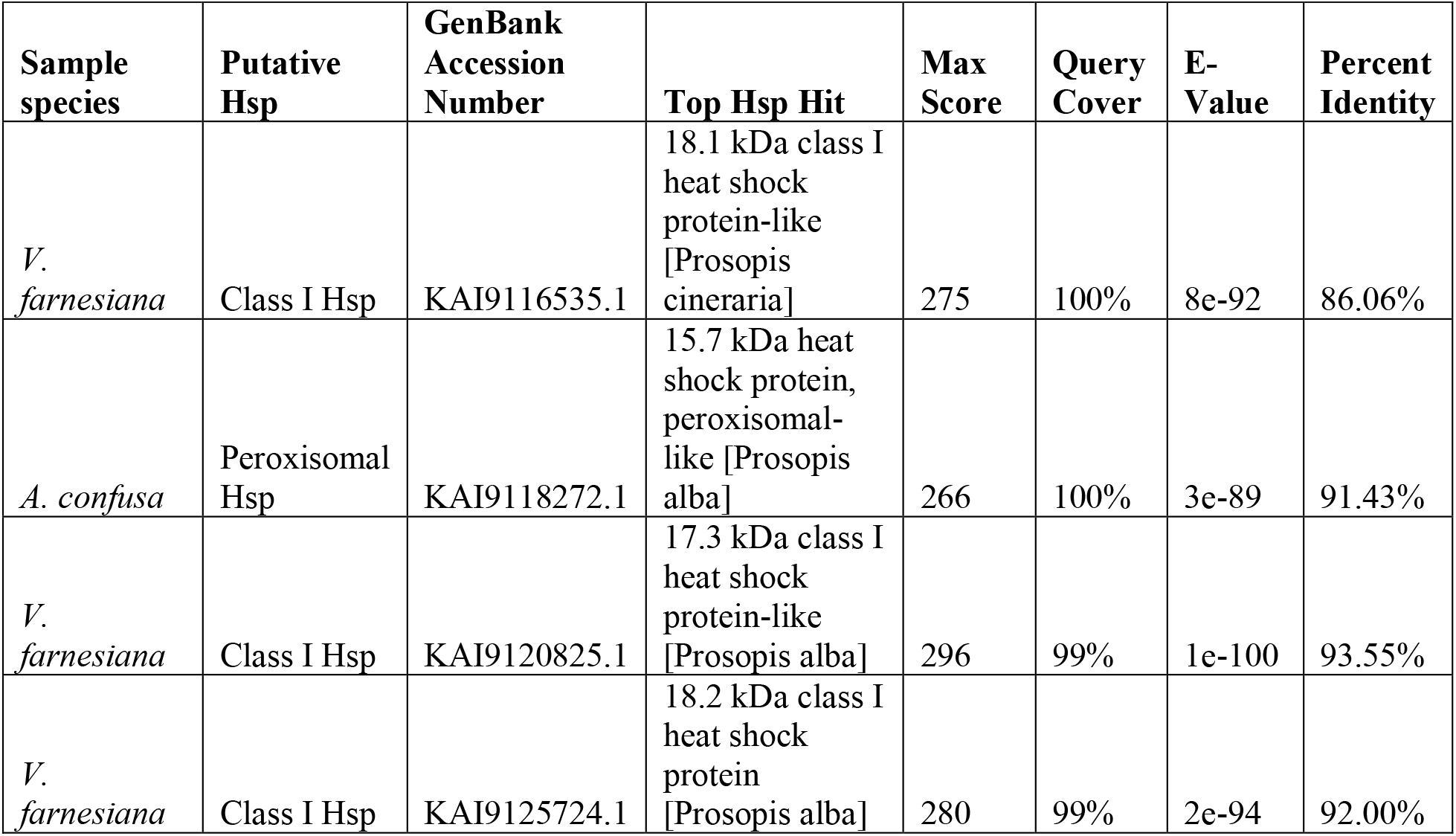
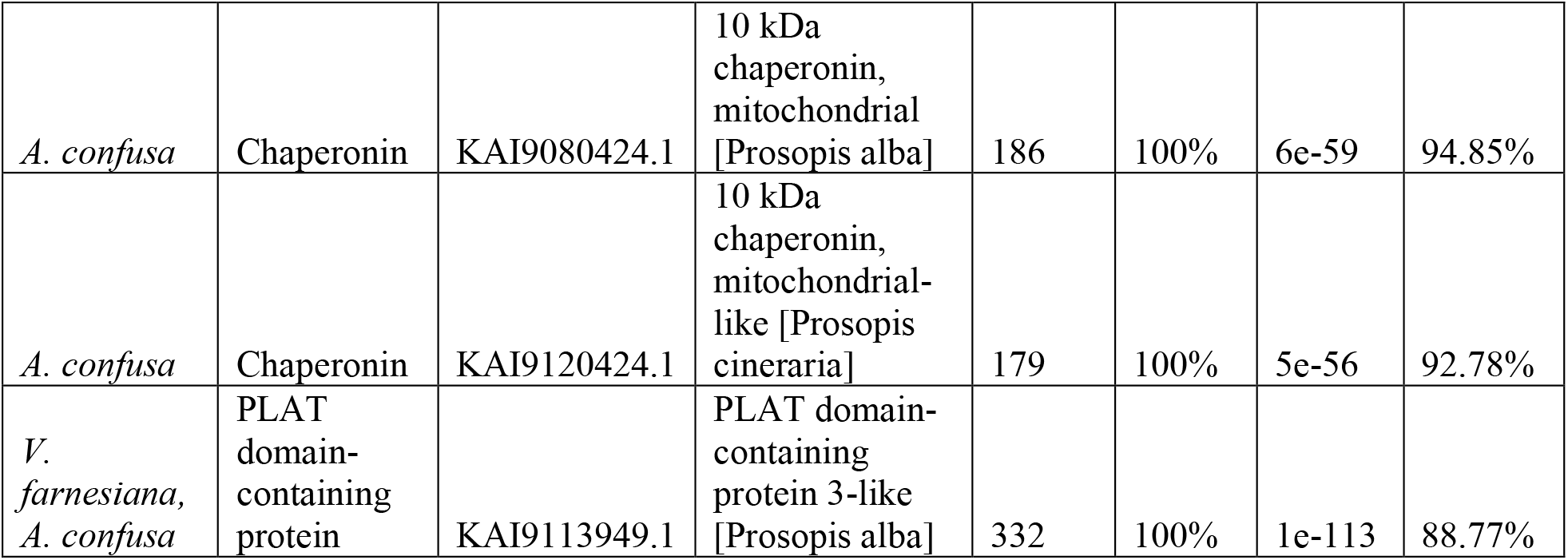
Plant small heat shock proteins identified in *A. confusa* and *V. farnesiana* nodule protein extracts, characterized using HPLC-MS peptide sequencing and BLAST comparison across legumes.

All rhizobial small heat shock proteins displayed total spectrum (as seen in Figure 1) counts greater than or equal to two in multiple samples (control and/or heat-treated bands), except heat shock protein IbpA which displayed 22 spectra counts in the control *V. farnesiana* sample and none in the heat-treated group.

**Figure 1:**
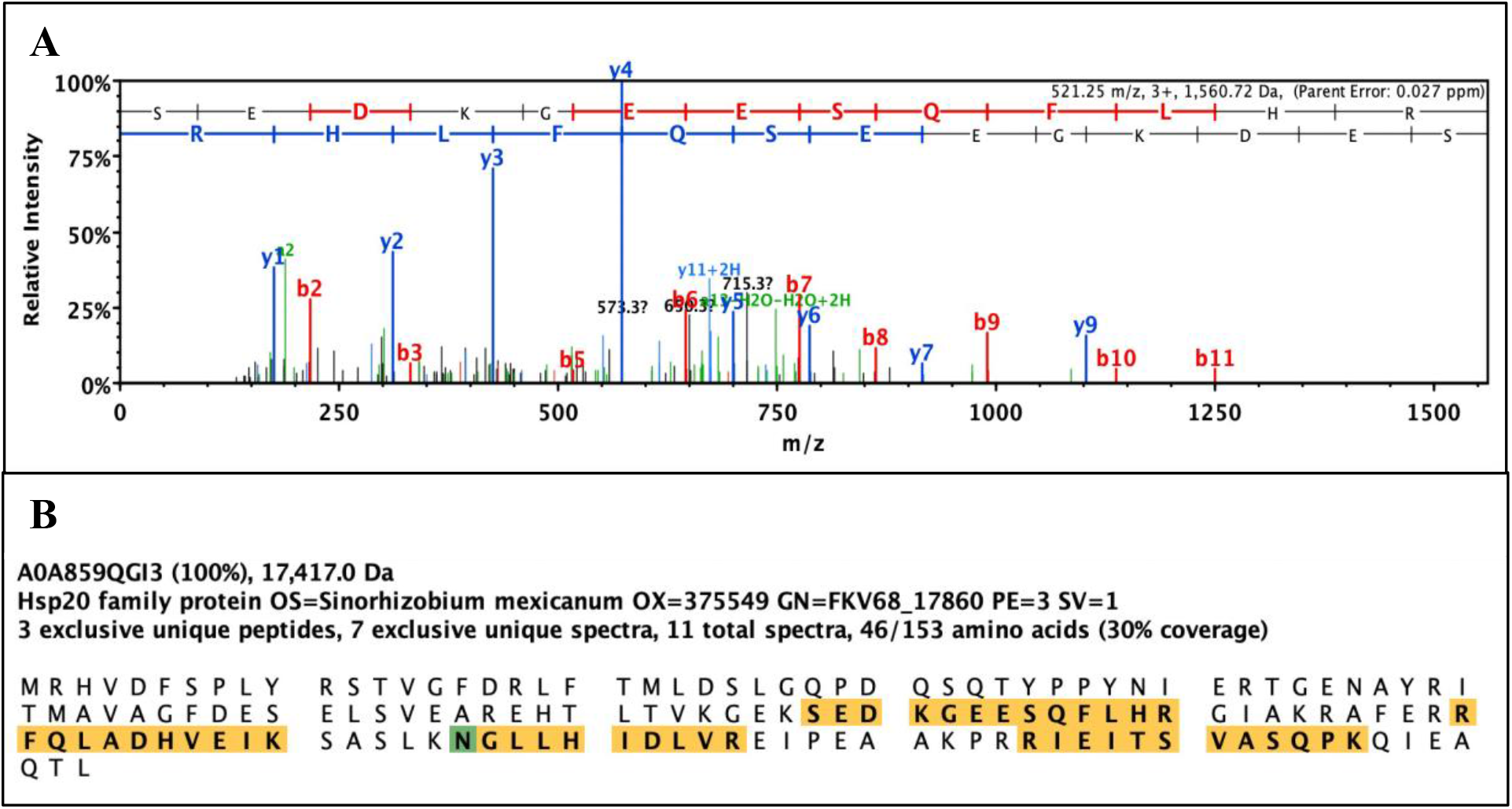
Example of peptide fragment spectral overlay and sequence alignment with Hsp20 family protein from *Sinorhizobium mexicanum*. Four bands were excised from two SDS-PAGE polyacrylamide gels (two bands each, one set around 18 kDa, the other around 16 kDa), one control band with protein content from non-treated excised nodules and one from resolved heat-treated nodule protein isolates. A. Overlay of HPLC-MS spectra from peptide fragments corresponding to rhizobial Hsp20 protein identified in an excised SDS-PAGE band. B. Sequence alignment of peptide fragments identified in HPLC-MS with reference full-protein sequence from UniProtKB (Accession number: A0A859QGI3).

Although Hsp20s were observed in control samples, which was unexpected given these proteins are not thought to be constitutively expressed, we observed higher expression of all Hsp20 variants (Hsp20 protein, Hsp20 family protein SHSP-domain proteins and small heat shock protein) in heat-treated samples relative to controls, as seen in the fold change values greater than two for all protein hits (Figure 2). Conversely, the chaperonin GroES hits displayed fold changes less than one, indicating this group of molecular chaperones is less inducible in these tissues, or is constitutively expressed by bacterial symbionts in nodule tissues whereas Hsp20 variants are more responsive to temperature and stress conditions.

**Figure 2:**
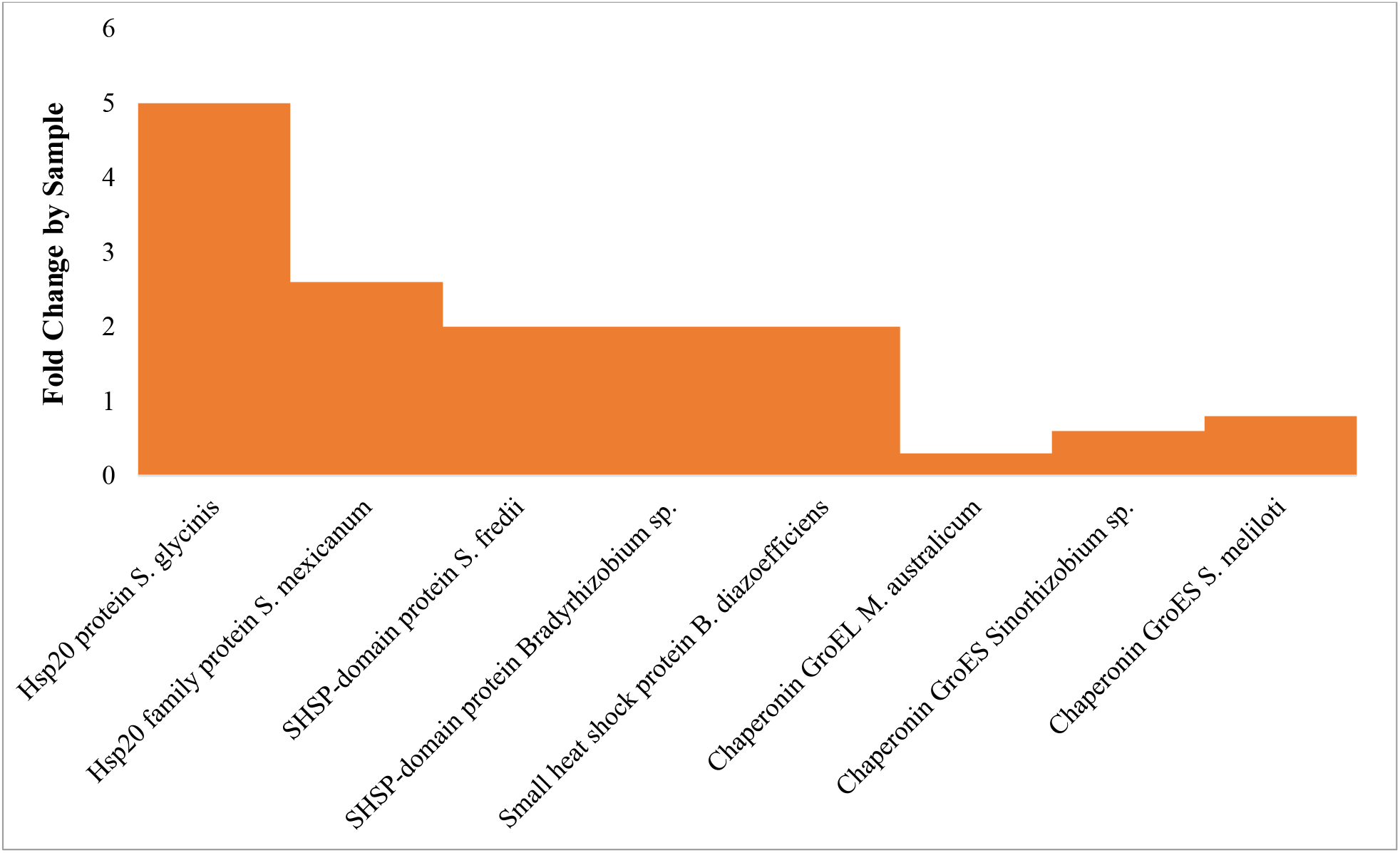
Rhizobial heat shock protein expression levels in heat-treated nodules relative to control nodules quantified by fold change in total spectrum count. Identified rhizobial Hsp20 (Table 1) expression was quantified by fold change in total spectrum count using Scaffold Q+ Scaffold_5.0.1 (Proteome Sciences) qualitative analysis function, first excluding all *A. confusa* samples and determining the fold change in heat-treated *V. farnesiana* hits relative to the control sample. Fold change in total spectrum counts of *A. confusa* heat-treated samples were then calculated relative to the control, excluding *V. farnesiana*. Heat shock protein IbpA excluded due to lack of identified total spectra in heat-treated sample.

There was no remarkable pattern observed in the fold change by sample of the plant sHsp hits. Two of the seven identified proteins (18.1 kDa class I Hsp and 10 kDa chaperonin KAI9113949.1) fold change could not be calculated as there were no hits observed in the respective *V. farnesiana* and *A. confusa* control samples. Of the remaining five identified plant sHsps two proteins (peroxisomal Hsp and 18.2 kDa class I Hsp) displayed fold changes exceeding two, while the other three exhibited a fold change near one.

### *Acacia confusa* root tissues express Hsp20 as observed with western blotting

To further investigate relative Hsp20 expression across species we conducted western blot experiments with available Hsp20 antibodies. Three experiments were conducted over six weeks, following the methodology outlined for the western blot experiments, with previously undisturbed individuals (*A. confusa* plants). Figure 3 displays the results of chemiluminescent imaging for all three western blot experiments. Each sample is labeled with tissue type (root or nodule) and treatment type (total control, stress control, heat-treated) corresponding to the treatment groups described in methods (heat shock induction of nodule/root tissues).

**Figure 3:**
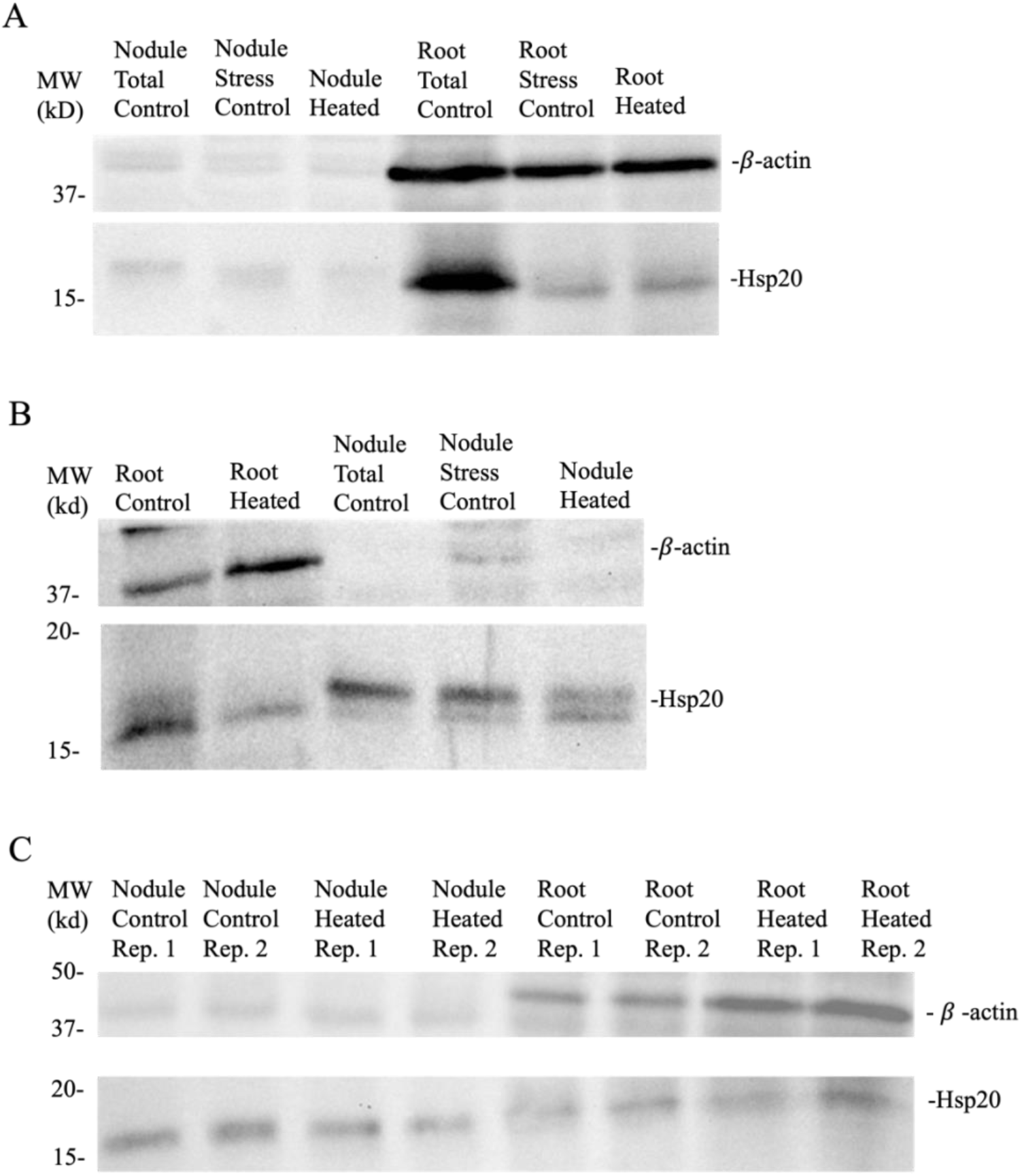
Chemiluminescent imaging of western blots with beta-actin and Hsp20 antibodies qualitatively validates Hsp20 expression observed in nodules and root protein extracts. A.) Western blot from the first experiment including all controls for both tissue types and beta-actin loading control; lanes 1-3 were cropped from the image to exclude data not presented. B.) Second western blot experiment excluding root tissue stress control (no root tissue harvested from stress control plant) including beta-actin loading control. C.) Third western blot experiment excluding stress control individual entirely, no stress control plant incorporated, with technical duplicates from the same individual (heat-treated or total control) that received separate chemical extractions (same methodology). Nodule protein extracts from *A. confusa* nodules, as well as *A. confusa* root tissue are imaged alongside a protein ladder (L). Inequal signal intensity of beta-actin, the loading control is evident, as well as qualitatively uniform expression of Hsp20 across control and heat-treated samples in all experiments. Images were inverted for improved contrast and band analysis using Fiji^17^. Data not presented in this paper has been removed from this figure, but it is otherwise unaltered (no change in contrast/brightness/etc.) Full gel images available in supplementary materials (Figures S2-S4).

We did not see standard beta-actin expression across tissue types, as one would expect for a loading control, which prevented us from conducting quantitative analysis of Hsp expression from the western blot signal intensities. In *A. confusa* root tissue, high beta-actin chemiluminescent signal intensity was observed, however near-zero signal was observed in *A. confusa* nodule samples. Hsp20 appeared to remain approximately consistent across control and heat-treated groups in both tissue types, although we are unable to conclude with certainty any patterns pertaining to Hsp expression from these experiments.

## Discussion

Our results validate that Hsp20 is expressed in nodule tissues; however, further research that quantifies relative Hsp20 expression across species occupying different ranges, and plant tissues is required to expand our understanding of the heat shock response in plants and its potential as a mechanism for thermoacclimation. It is interesting to note that both plant and rhizobial sHsps were identified in both *V. farnesiana* and *A. confusa* nodule protein isolates, potentially indicating that the HSR in nodule tissues is influenced and carried out by both bacterial and plant cells in nodule tissues.

Of note, we observed a wide variety of *A. pycnatha* sHsps including the mitochondrial chaperonin 10 and a PLAT domain-containing protein observed in both sample species in our experiment. Plants have been observed to express proteins such as Hsp10, or mitochondrial chaperonins that act as co-chaperones to aid in the function of Hsp60s^18^. Chaperonin, or Hsp10, has been identified in a variety of plant species in a variety of plant tissues^18^, and from these results is likely expressed in *A. confusa* nodule tissues as well. The PLAT (Polycystin-1, Lipoxygenase, Alpha Toxin) domain associates with lipids, and many PLAT domain-containing proteins have been linked to abiotic stress, including temperature, salt, and drought stress^19^. The PLAT domain is thought to play a role in protein-protein interactions and protein-membrane regulation and has been implicated in catalytic regulation and substrate specificity of its enzyme binding partners that effectively promotes growth during stress conditions^20^. The presence of the PLAT domain-containing protein in our analysis is interesting to note, although fold change in expression of this protein was variable across the two species sample and it is difficult to discern if this protein may play a role in the HSR, or if another stress condition experienced by plants grown in our greenhouse setup triggered expression of these stress proteins.

The difference in fold change exhibited by rhizobial Hsp20s compared to that of the GroES chaperone species is an interesting observation as it may indicate that these distinct groups of Hsps could function differently in terms of inducibility. Where rhizobial Hsp20s displayed substantial increase in expression, as quantified by total spectrum count, after heat-treatment, the GroES chaperones displayed no such change, potentially indicating that these Hsps are constitutively expressed by rhizobia in nodules or are less inducible than their Hsp20 counterparts. These results are unexpected given GroES, also known as a Hsp10 chaperone protein common in bacteria, is well-studied and it has been observed to be overexpressed during heat-stress and it thought to be inducible with moderate heat stress conditions^21^.

Unfortunately, we were unable to address our second research aim as inconsistent beta-actin expression prevented qualitative analysis that might reveal patterns in Hsp20 expression across tissue types. However, we can conclude that Hsp20s are produced in both nodule and root tissues due to the qualitative observance of chemiluminescent signals in almost sample types, suggesting some level of Hsp20 expression has occurred. Our choice to use beta-actin as a loading control was ineffectual, as rhizobia are not known to express beta-actin and although it has been observed that plant cells do contain beta-actin during nodule development (and after nodule formation)^22,23^, it is not surprising that we observed much higher beta-actin signal intensity in root tissue relative to the nodule protein extracts, as these tissues vary in their density of plant cells and therefore will likely vary in their beta-actin content. Given the extreme difference in symbiont and host proteome composition and in light of recent criticism of ‘house-keeping gene’ loading controls, it would be interesting to attempt these experiments using the Total Protein Analysis method for signal intensity standardization^24^. Normalizing the Hsp20 signal intensities to total protein content, quantified by a colorimetric stain intensity, would likely allow one to compare Hsp20 expression across tissue types with greater confidence^24^. Previous studies have found that responses to drought stress induce different proteomic effects in nodule bacteroids relative to plant tissues^25^, thus it is not outlandish to suggest that there may indeed be differential Hsp20 expression in one tissue type relative to the other, however our data is not sufficient to validate a statistically significant quantitative relationship. Moreover, we observed Hsp20 signal in almost all control samples, which is unideal given Hsps, including Hsp20 are canonically stress-induced, thus we theorize that Hsp20 may be constitutively expressed in both tissue types of *A. confusa* under the specific greenhouse conditions of our experiments. Future experiments should seek to design control conditions that more appropriately establish baseline Hsp expression, or to examine the effect of artificial greenhouse conditions on cellular stress processes in these plant tissues.

Data not presented in this article included western blot results of *Robinia pseudoacacia* protein extracts, which exhibited highly variable beta-actin and Hsp20 expression as observed by highly variable chemiluminescent signal intensities. The incorporation of *Robinia* samples was in an attempt to discern potential variability of Hsp20 expression in nodules and/or root tissues of temperate N-fixer trees relative to their tropical counterparts. We struggled to culture healthy *Robinia* plants in our greenhouse setting as light and temperature conditions were maintained at levels that could sustain tropical tree species and may have led to heat-stress in our temperate species before heat shock induction which we postulate was the driver of our inconsistent results. Future experiments should aim to appropriately compare temperate versus tropical Hsp20 expression as variation in biochemical responses to heat stress may very well be a driver of N-fixation regulation on larger scales, but we were unable to produce data to support such claims. As climate change continues to alter global cycles that are integral for ecosystem stability and agriculture, we should continue to research mechanisms of adaptation that may influence our projections of global change impacts and inform environmental policy worldwide.

## Acknowledgments

We would like to acknowledge the Proteomics and Molecular Analysis Platform at the RI-MUHC for their work in processing samples with HPLC-MS, performing sequence analysis with reference databases, and producing scaffold files for nodule proteome analysis.

## Data Availability

The mass spectrometry proteomics data have been deposited to the ProteomeXchange Consortium via the PRIDE partner repository with the dataset identifier PXD055599.

## Author contributions

CGF, RF and FMS conceived and designed the study. CGF and RF performed the research. CGF analyzed proteomic data. All authors interpreted results and contributed to the manuscript - writing led by CGF and FMS.

## References

1. Menge, D. N. L.; Lichstein, J. W.; Ángeles-Pérez, G. Nitrogen Fixation Strategies Can Explain the Latitudinal Shift in Nitrogen-Fixing Tree Abundance. Ecology 2014, 95 (8), 2236–2245.

2. Bytnerowicz, T. A.; Min, E.; Griffin, K. L.; Menge, D. N. L. Repeatable, Continuous and Real-time Estimates of Coupled Nitrogenase Activity and Carbon Exchange at the Whole-plant Scale. Methods Ecol Evol 2019, 10 (7), 960–970.

3. Udvardi, M.; Poole, P. S. Transport and Metabolism in Legume-Rhizobia Symbioses. Annu. Rev. Plant Biol. 2013, 64 (1), 781–805.

4. Lindström, K.; Mousavi, S. A. Effectiveness of Nitrogen Fixation in Rhizobia. Microbial Biotechnology 2020, 13 (5), 1314–1335.

5. Bytnerowicz, T. A.; Akana, P. R.; Griffin, K. L.; Menge, D. N. L. Temperature Sensitivity of Woody Nitrogen Fixation across Species and Growing Temperatures. Nat. Plants 2022, 8 (3), 209–216.

6. Houlton, B. Z., Wang, Y.-P., Vitousek, P. M. & Field, C. B. A unifying framework for dinitrogen fixation in the terrestrial biosphere. Nature 2008, 454, 327–330.

7. Wahid, A.; Gelani, S.; Ashraf, M.; Foolad, M. Heat Tolerance in Plants: An Overview. Environmental and Experimental Botany 2007, 61 (3), 199–223.

8. Al-Whaibi, M. H. Plant Heat-Shock Proteins: A Mini Review. Journal of King Saud University - Science 2011, 23 (2), 139–150.

9. Yang, Z.; Du, H.; Xing, X.; Li, W.; Kong, Y.; Li, X.; Zhang, C. A Small Heat Shock Protein, GmHSP17.9, from Nodule Confers Symbiotic Nitrogen Fixation and Seed Yield in Soybean. Plant Biotechnology Journal 2022, 20 (1), 103–115.

10. Reich, B. P.; Wright, I. J.; Cavender-Bares, J.; Craine, J. M.; Oleksyn, J.; Westoby, M.; Walters, M. B. The Evolution of Plant Functional Variation: Traits, Spectra, and Strategies. Int. J. Plant Sci. 2003, 164(S3), S143–S164.

11. Frova, C.; Gorla, M. S. Quantitative Expression of Maize HSPs: Genetic Dissection and Association with Thermotolerance. Theoret. Appl. Genetics 1993, 86–86 (2–3), 213–220.

12. Wang, W.; Vignani, R.; Scali, M.; Cresti, M. A Universal and Rapid Protocol for Protein Extraction from Recalcitrant Plant Tissues for Proteomic Analysis. Electrophoresis 2006, 27 (13), 2782–2786.

13. Rodrigues, E. P.; Torres, A. R.; da Silva Batista, J. S.; Huergo, L.; Hungria, M. A Simple, Economical and Reproducible Protein Extraction Protocol for Proteomics Studies of Soybean Roots. Genet. Mol. Biol. 2012, 35 (1 Suppl), 348–352.

14. Kruger, N. J. The Bradford Method for Protein Quantitation. The Protein Protocols Handbook; Walker, J. M., Ed.; Springer Protocols Handbooks; Humana Press: Totowa, NJ, 2009; pp 17–24.

15. Weber, K.; Osborn, M. The reliability of molecular weight determinations by dodecyl sulfate-polyacrylamide gel electrophoresis. J. Biol. Chem. 1969, 244 (16), 4406–4412.

16. McLay, T.; Murphy, D.; Holmes, G.; Mathews, S.; Brown, G. et al. A genome resource for Acacia, Australia’s largest plant genus. PLoS one 2022, 17(10), e0274267.

17. Schindelin, J., Arganda-Carreras, I., Frise, E., Kaynig, V., Longair, M., Pietzsch, T., et al. Fiji: an open-source platform for biological-image analysis. Nature Methods 2012, 9(7), 676–682.

18. Pareek, A.; Mishra, D.; Rathi, D.; Verma, J. K.; Chakraborty, S.; Chakraborty, N. The small heat shock proteins, chaperonin 10, in plants: An evolutionary view and emerging functional diversity. J. Exp. Bot. 2021, 182, 104323.

19. Trujillo, D. I.; Silverstein, K.; Young, N. Nodule-specific PLAT domain proteins are expanded in the Medicago lineage and required for nodulation. New Phytologist 2019, 222 (3), 1538–1550.

20. Hyun, T. K.; van der Graaff, E.; Albacete, A.; Eom, S. H.; Großkinsky, D.; Böhm, H.; Janschek, U.; Rim, Y.; Ali, W.; Kim, S. Y.; Roitsch, T. The Arabidopsis PLAT Domain Protein1 Is Critically Involved in Abiotic Stress Tolerance. PLoS one 2014, 9 (11), e112946.

21. Ventura, M.; Canchaya, C.; Zink, R.; Fitzgerald, G.; van Sinderen, D. Characterization of the groEL and groES Loci in Bifidobacterium breve UCC 2003: Genetic, Transcriptional, and Phylogenetic Analyses. Appl. Environ. Microbiol. 2004, 70 (10), 6197–6209.

22. Zhang, X.; Han, L.; Wang, Q.; Zhang, C.; Yu, Y.; Tian, J.; Kong, Z. The host actin cytoskeleton channels rhizobia release and facilitates symbiosome accommodation during nodulation in Medicago truncatula. New Phytologist 2019, 211, 1049–1059.

23. Yokota, K.; Fukai, E.; Madsen, L.; Jurkiewicz, A.; Rueda, P. et al. Rearrangement of Actin Cytoskeleton Mediates Invasion of Lotus japonicus Roots by Mesorhizobium loti. The Plant Cell 2009, 21 (1), 267–284.

24. Eaton, S.; Roche, S.; Hurtado, M.; Oldknow, K.; Farquharson, C.; Gillingwater, T.; Wishart, T. Total Protein Analysis as a Reliable Loading Control for Quantitative Fluorescent Western Blotting. PLoS one 2013, 8 (8), e72457.

25. Larrainzar, E.; Wienkoop, S.; Weckwerth, W.; Ladrera, R.; Arrese-Igor, C.; González, E. Medicago truncatula Root Nodule Proteome Analysis Reveals Differential Plant and Bacteroid Responses to Drought Stress. Plant Physiol. 2007, 144 (3), 1495–1507.

