## Supplementary Materials for "Small heat shock proteins (sHSPs) identified in nodules of tropical woody legumes"

**Supplementary Material:**

**Table S1: Inoculum Bradyrhizobia and Characterized Associate *Acacia* Species for Nodule Development and Nitrogen Fixation.**

| **Bradyrhizobium strain USDA code** | **Associate Legume** |
| --- | --- |
| 3001 | *Acacia decurrens* |
| 3003 | *Acacia linifolia* |
| 3328 | *Acacia pennatula* |
| 3475 | *Acacia melanoxylon* |
| 3517 | *Acacia albida* |
| 3841 | *Acacia constricta* |
| 4361 | *Acacia angustissima* |
| 4400 | *Acacia senegal* |
| 4869 | *Acacia senegal* |
| HM2 | Unlisted |
| HM3 | Unlisted |

**Table S2: Modified Hoagland’s Fertilizer Recipe for Inducing Nitrogen Limitation Across Species.**

| **Hoagland’s Ingredient** | **Concentration (mL/L)** |
| --- | --- |
| 0.5M CaCl_2_ | 2.25 |
| 1M Ca(NO_3_)_2_ | 2.25 |
| 1M KCl | 1.5 |
| 1M MgSO_4_ | 0.6 |
| Micronutrients (H_3_BO_3_, MnCl_2_, ZnCl_2_, CuCl_2_, and Na_2_MoO_4_) | 0.3 |
| Fe-EDTA | 0.3 |
| KH_2_PO_4_ | 0.3 |

**Figure S1: ClustalOmega Multiple Sequence Alignment of Hsp20 family protein, *Sinorhizobium mexicanum* with HspB6 of Human, Mouse, and Rat**
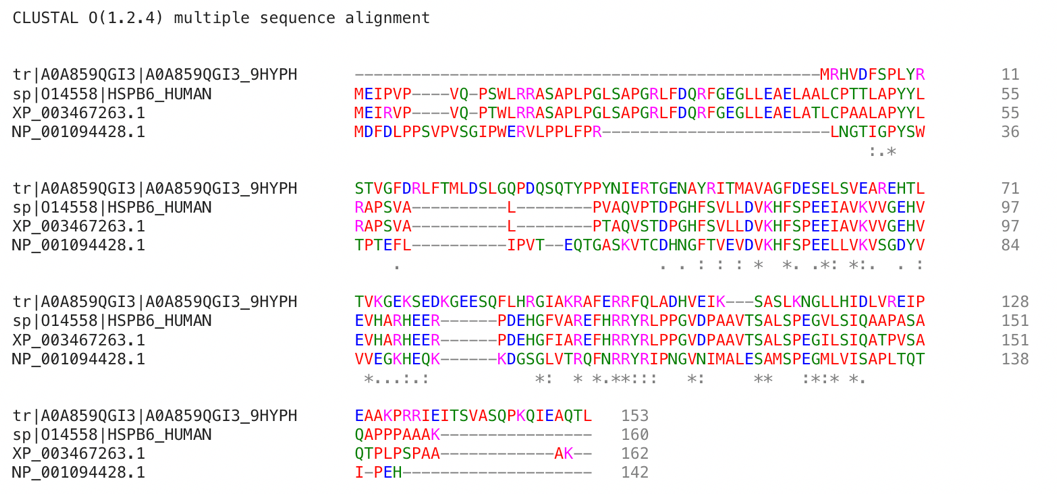

**Figure S2: Unannotated Western Blot 1 with *Acacia confusa* Nodule and Root Protein Extracts**

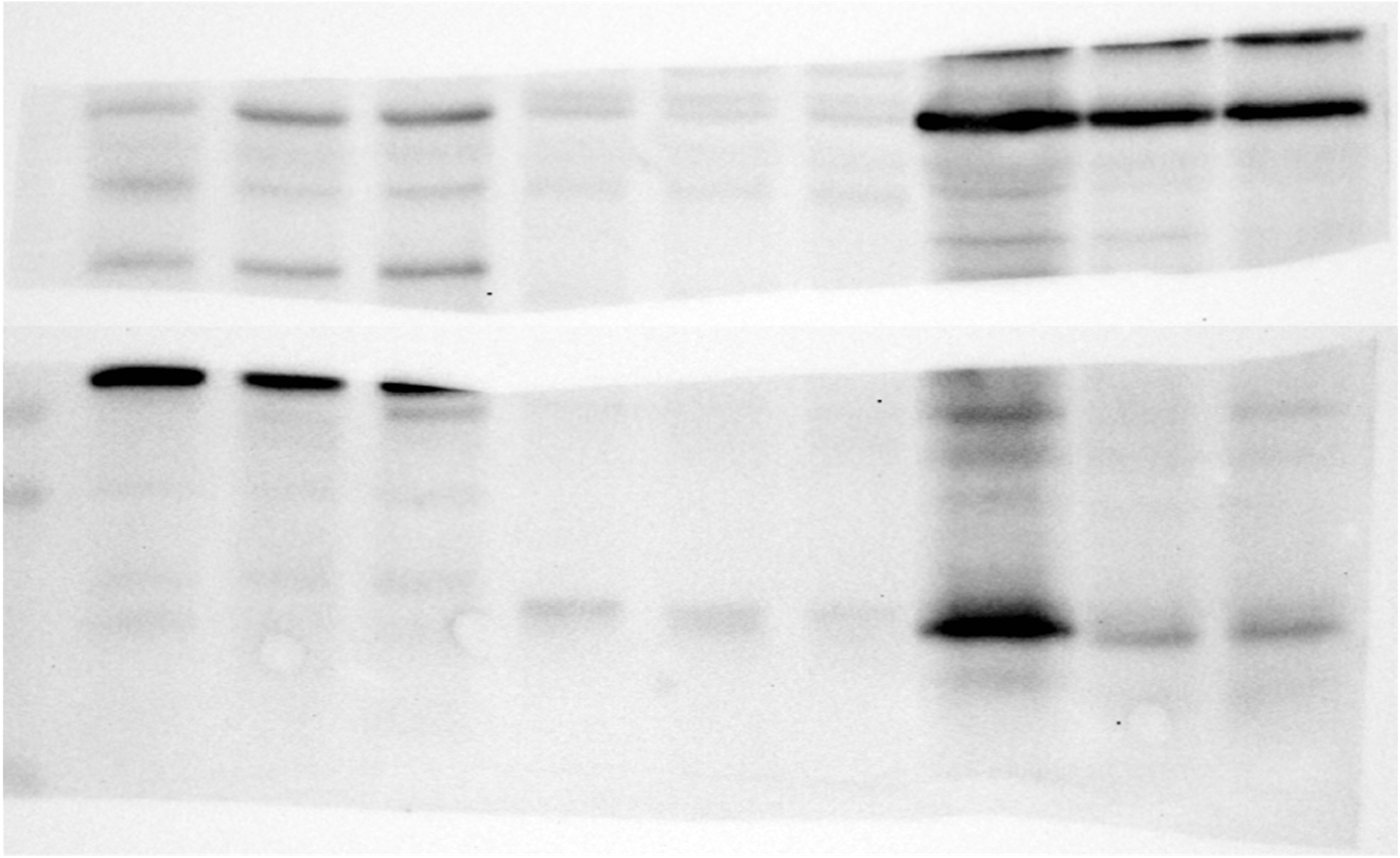

**Figure S3: Unannotated Western Blot 2 with *Acacia confusa* Nodule and Root Protein Extracts**

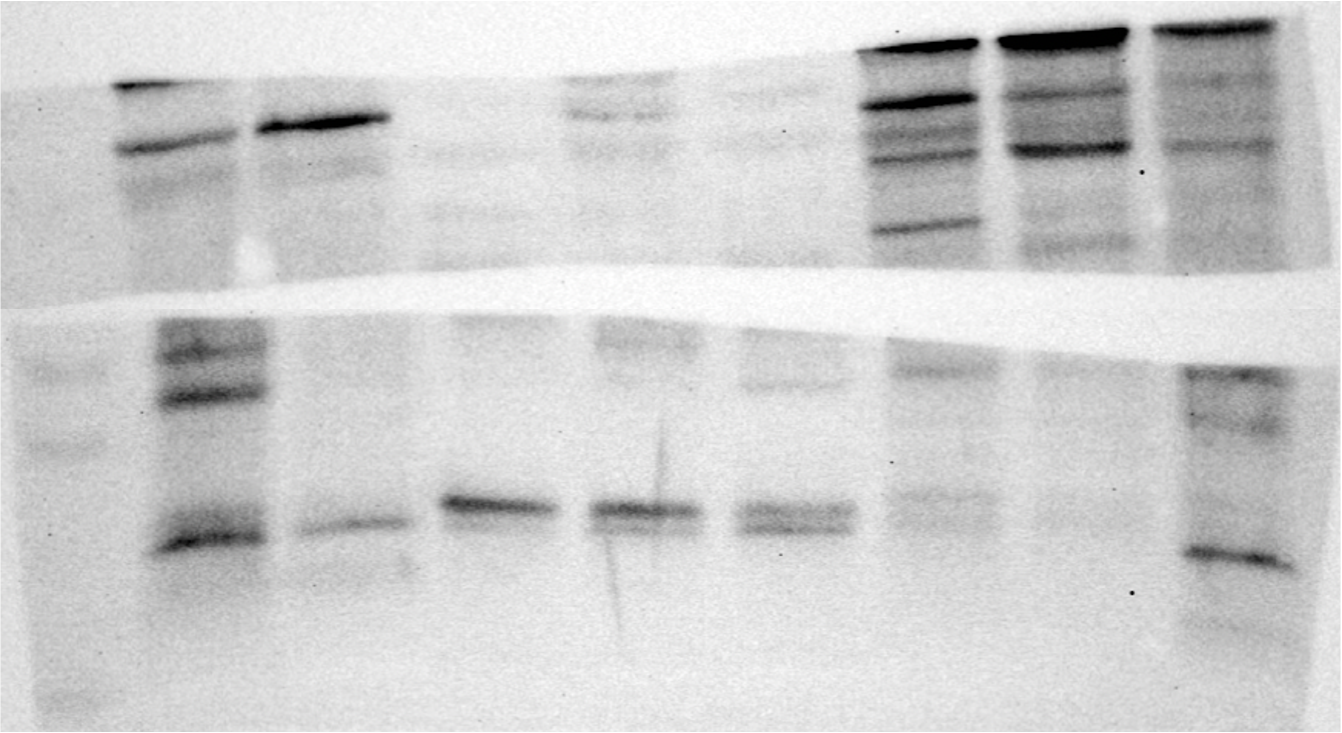

**Figure S4: Unannotated Western Blot 3 with *Acacia confusa* Nodule and Root Protein Extracts**

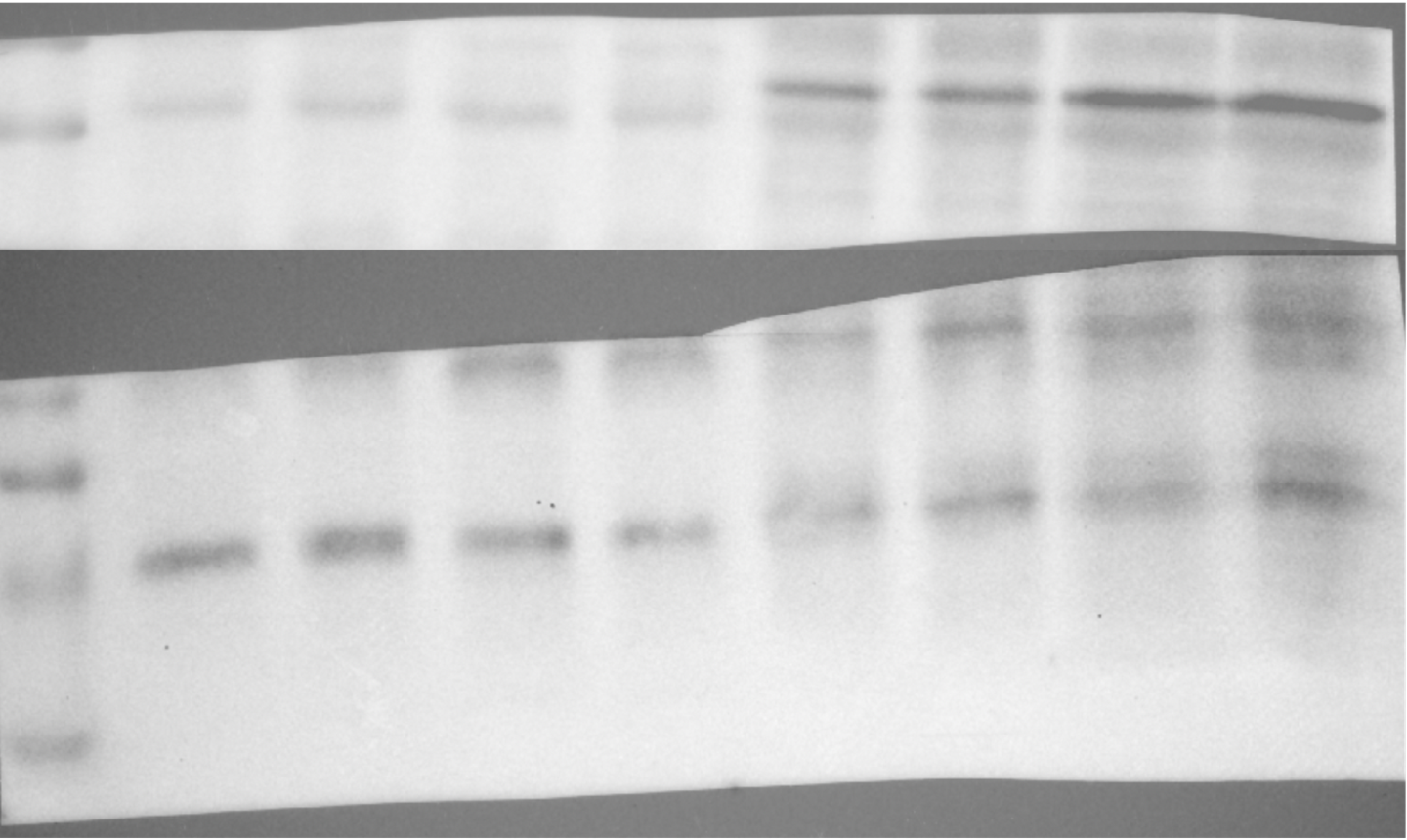
